# High-order functional interactions in ageing explained via alterations in the connectome in a whole-brain model

**DOI:** 10.1101/2021.09.15.460435

**Authors:** Marilyn Gatica, Fernando E. Rosas, Pedro A.M. Mediano, Ibai Diez, Stephan P. Swinnen, Patricio Orio, Rodrigo Cofré, Jesus M. Cortes

**Affiliations:** Centro Interdisciplinario de Neurociencia de Valparaíso, Universidad de Valparaíso, Valparaíso, Chile; Biomedical Research Doctorate Program, University of the Basque Country (UPV/EHU), 48940 Leioa, Spain; Department of Psychology, University of Cambridge, Cambridge, UK; Centre for Psychedelic Research, Department of Brain Science, Imperial College London, London SW7 2DD, UK; Data Science Institute, Imperial College London, London SW7 2AZ, UK; Department of Radiology, Division of Nuclear Medicine and Molecular Imaging, Massachusetts General Hospital and Harvard Medical School. Boston, USA; Gordon Center for Medical Imaging, Department of Radiology, Massachusetts General Hospital and Harvard Medical School. Boston, USA; Athinoula A. Martinos Center for Biomedical Imaging, Massachusetts General Hospital, Harvard Medical School. Boston, USA; Research Center for Movement Control and Neuroplasticity, Department of Movement Sciences, KU Leuven, Leuven, Belgium; KU Leuven Brain Institute (LBI), KU Leuven, Leuven, Belgium; Instituto de Neurociencias, Facultad de Ciencias, Universidad de Valparaíso, Valparaíso, Chile; CIMFAV-Ingemat, Facultad de Ingeniería, Universidad de Valparaíso, Valparaíso, Chile; Department of Integrative and Computational Neuroscience, Paris-Saclay Institute of Neuroscience, Centre National de la Recherche Scientifique, Gif-sur-Yvette, France; Computational Neuroimaging Lab, Biocruces-Bizkaia Health Research Institute, 48903 Barakaldo, Spain; IKERBASQUE: The Basque Foundation for Science, 48013 Bilbao, Spain; Department of Cell Biology and Histology, University of the Basque Country, 48940 Leioa, Spain

## Abstract

The human brain generates a rich repertoire of spatio-temporal activity patterns, which support a wide variety of motor and cognitive functions. These patterns of activity change with age in a multi-factorial manner. One of these factors is the variations in the brain’s connectomics that occurs along the lifespan. However, the precise relationship between high-order functional interactions and connnectomics, as well as their variations with age are largely unknown, in part due to the absence of mechanistic models that can efficiently map brain connnectomics to functional connectivity in aging. To investigate this issue, we have built a neurobiologically-realistic whole-brain computational model using both anatomical and functional MRI data from 161 participants ranging from 10 to 80 years old. We show that the age differences in high-order functional interactions can be largely explained by variations in the connectome. Based on this finding, we propose a simple neurodegeneration model that is representative of normal physiological aging. As such, when applied to connectomes of young participant it reproduces the age-variations that occur in the high-order structure of the functional data. Overall, these results begin to disentangle the mechanisms by which structural changes in the connectome lead to functional differences in the ageing brain. Our model can also serve as a starting point for modelling more complex forms of pathological ageing or cognitive deficits.

**Author summary:** Modern neuroimaging techniques allow us to study how the human brain’s anatomical architecture (a.k.a. structural connectome) changes under different conditions or interventions. Recently, using functional neuroimaging data, we have shown that complex patterns of interactions between brain areas change along the lifespan, exhibiting increased redundant interactions in the older population. However, the mechanisms that underlie these functional differences are still unclear. Here, we extended this work and hypothesized that the variations of functional patterns can be explained by the dynamics of the brain’s anatomical networks, which are known to degenerate as we age. To test this hypothesis, we implemented a whole-brain model of neuronal activity, where different brain regions are anatomically wired using real connectomes from 161 participants with ages ranging from 10 to 80 years old. Analyzing different functional aspects of brain activity when varying the empirical connectomes, we show that the increased redundancy found in the older group can indeed be explained by precise rules affecting anatomical connectivity, thus emphasizing the critical role that the brain connectome plays for shaping complex functional interactions and the efficiency in the global communication of the human brain.

## Introduction

Advancing the neuroscience of ageing is highly relevant from a socio-economic perspective as the population of older adults is rising dramatically worldwide, with predictions foreseeing the percentage of people aged 60 years or older to increase from 900 million in 2015 up to 2.1 billion by 2050 [1]. An important challenge of contemporary neuroscience is to better understand the systemic effects of ageing on the structure and function of the human brain [2–5]. Ageing is a major risk factor for late-onset brain disorders, accelerates cognitive and motor decline and worsens the quality of life. A deeper understanding of this process could motivate novel interventions or protection therapies against age-related deterioration or neurodegenerative diseases [6–10].

Several effects of ageing on brain structure have been previously studied. Along the lifespan, the total brain volume increases from childhood to adolescence by approximately 25% on average, remains constant for the three following decades, and finally decays back to childhood size at late ages [11]. Additionally, it has been shown that the amount of atrophy in aged brains is not homogeneous, but some anatomical regions are more affected than others — well-known ageing-targeted structures are the hippocampus [12], prefrontal cortex [13] and basal ganglia [14,15]. White-matter degenerates faster than gray-matter along the lifespan, indicating that the overall connectivity is diminished with age [16]. Moreover, a progressive decrease in many tract-integrity measures has been shown using diffusion imaging, which is more pronounced in subjects above 60 years of age [5,6].

Previous studies of functional connectivity along the lifespan during resting state have shown that regions within the default mode network (DMN) become less functionally connected with age [4,7,17,18]. Additionally, the frontoparietal, dorsal attention, and salience networks also show some degree of age-related decline, including reduced within-network connectivity [17,19–21]. In contrast, between-network connectivity increase with age between the DMN, somatosensory, and the frontoparietal control networks. A stronger connectivity between the frontoparietal and dorsal attention networks has been reported as well [3,19,22,23]. Furthermore, recent investigations on high-order functional interactions have shown the predominance of redundant interdependencies in older adults, consistently across scales [24] and particularly relevant for the DMN [25].

Taken together, these findings suggest a general loss of functional specificity or circuit segregation across brain circuits [26]. Moreover, an increase in between-network interactions is a dominant feature of aging brains. Albeit important progress in our understanding of the effects of ageing on brain function, these effects are less understood than the effects on structural connectivity, which shows progressive age-related disconnection.

Despite these considerable advances in understanding how the anatomical and functional connectivity change along the lifespan, the relationship between changes in brain structure and function leading to age-related decline remains largely unknown. To bridge this important gap, we sought to investigate how age-related changes in brain structure affect its function. We tackle this fundamental question via whole-brain computational modelling, which is an emerging powerful tool to investigate the neurobiological mechanisms that underlie macroscale neural phenomena [27–29]. Our approach is based on the Dynamic Mean Field (DMF) model, which simulates mesoscale neural dynamics using coupled stochastic differential equations incorporating realistic aspects of neurophysiology [30–32]. DMF modelling can be used for systematically perturbing connectome characteristics while assessing the resulting effects on macroscale brain dynamics and function, thus opening the way to provide mechanistic modeling interpretations to data obtained from lesion studies and ageing [33–35].

Furthermore, the different DMF inputs and biophysical parameters can be systematically altered in ways that are beyond the capabilities of current experimental research, which makes whole-brain computational modelling a privileged tool to investigate the causal mechanisms that drive the brain’s organization and function [28,36–38].

Capitalising on this powerful approach, here we employed the DMF model informed by multimodal neuroimaging — including empirical BOLD fMRI dynamics and anatomical connectivity obtained from diffusion MRI. In particular, we used the DMF model to investigate how changes in the human connectome induced by ageing shape the changes in high-order brain interdependencies. Following the methodology put forward in Ref. [39], our analysis focuses on two types of qualitatively distinct high order interactions which are critical in our approach: synergy and redundancy. The latter is an extension of the standard notion of correlation that applies to three or more variables, in which each variable has a ‘copy’ of some common information shared with other variables [39]. An example of extreme redundancy is full synchronisation, where the value of one signal enables one to predict the state of any other. In contrast, synergy refers here to statistical relationships that regulate a group of variables but do not constrain any subset of them [40, 41]. Synergy enables a counterintuitive coexistence of local independency and global cohesion, proposed to be instrumental for high order brain functions, while redundancy — including cases of high synchronization such as deep sleep or epileptic seizures — would make neural systems less well-suited for them [42–45]. Our results show that DMF reproduces age-related changes in redundancy and synergy, previously reported in Ref. [24], and that these changes are driven by a process of connectome neurodegeneration. Leveraging this evidence, we developed a non-linear neurodegenerative model that efficiently mimics connectome ageing and its associated changes in the pattern of high-order interactions of the human brain.

## Results

### The DMF model reproduces empirical differences between age groups in the high-order redundant and synergistic interactions

Our analyses are based on the DMF model, which uses structural and functional connectivity matrices, (respectively SC and FC), to simulate the activity of various brain regions wired with SC in presence of local excitatory and inhibitory neuronal populations. A biophysical haemodynamic model [46] is then used to transform the DMF model’s firing rate dynamics into BOLD-like signals. The DMF is calibrated by optimising a free parameter, denoted by *G*, that allows the model to best approximate pairwise functional activity [28]. This procedure is illustrated in Fig. 1, and details are provided in methods.

**Fig 1.**
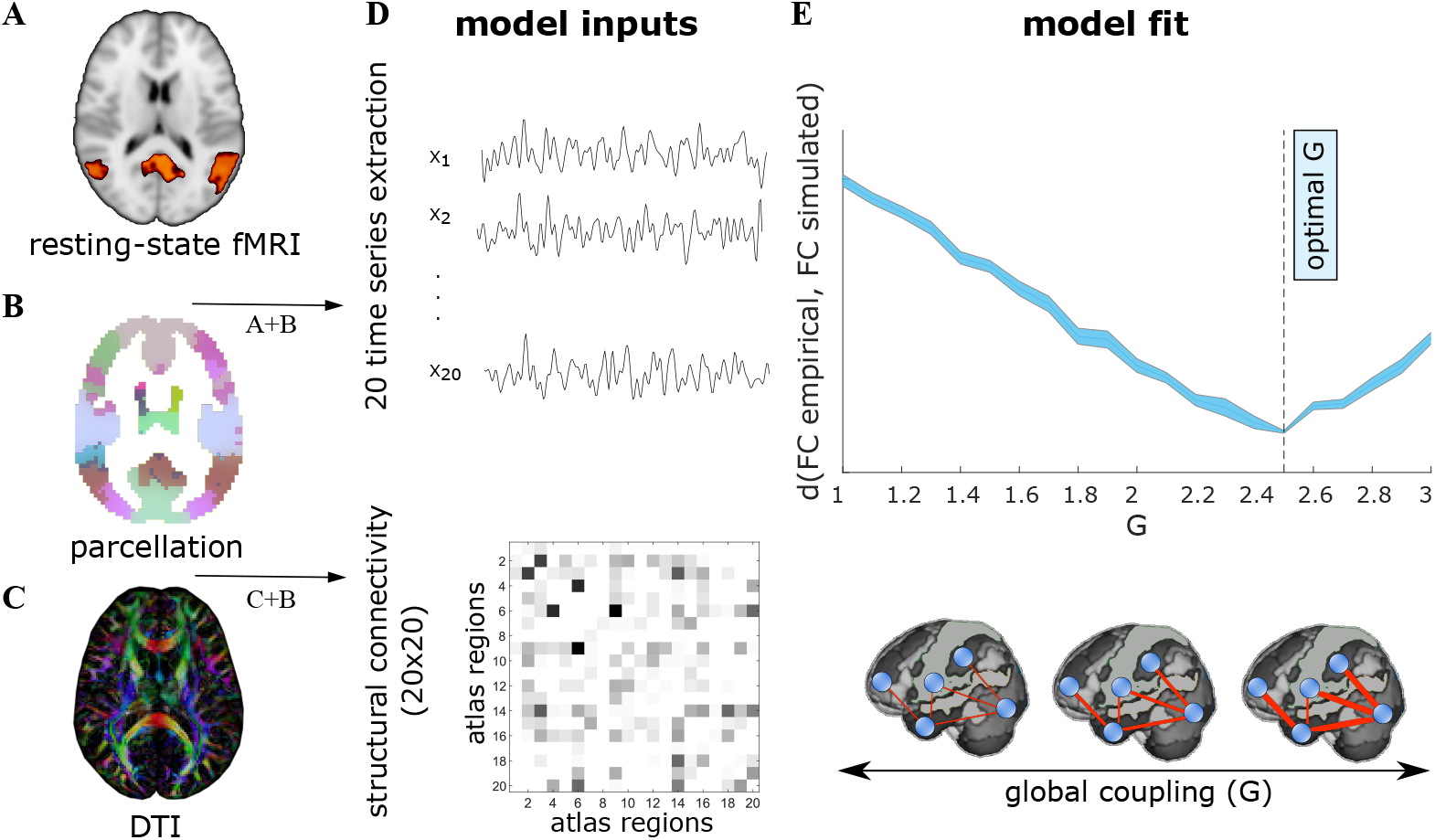
Whole-brain Dynamic Mean Field Model. The DMF model has several inputs: **A**: BOLD signals from fMRI data, **B**: a parcellation, in our case comprising 20 atlas regions, and **C**: the individual connectome obtained from DTI. **D**: Applying the parcellation to the fMRI and DTI datasets, we obtained 20 time-series signals and a 20 × 20 matrix representing the connectome. **E**: BOLD-like signals are simulated using the connectome and different values of the coupling parameter *G*. For each *G*, we compare the simulated and empirical data using the Kolmogorov-Smirnov distance (*d*) over the distribution of values of the FC matrices. Finally, the selected optimal *G* is the value that minimizes this distance.

To study the effect of ageing on brain dynamics, we used functional data from different participants divided into four age groups, similar to previous work [24]: I1 (*N*_1_ = 28 participants, age 10-20 years), I2 (*N*_2_ = 46, 20-40 years), I3 (*N*_3_ = 29, 40-60 years) and I4 (*N*_4_ = 58, 60-80 years). One DMF model was built for each age group, using the average SC within each group (Fig. 2A). Next, each model was calibrated separately, resulting in one *G* value per group (Fig. 2B). For each model, we simulated the brain activity using different random seeds, and the high-order interdependencies were calculated from these simulated data. In particular, we calculated the O-information [39], which can be considered an extension of the neural complexity previously proposed in [47] under the light of Partial Information Decomposition [48]. In essence, the O-information captures the balance between redundancies and synergies in a set of interacting variables [49,50] (for further details see Methods).

**Fig 2.**
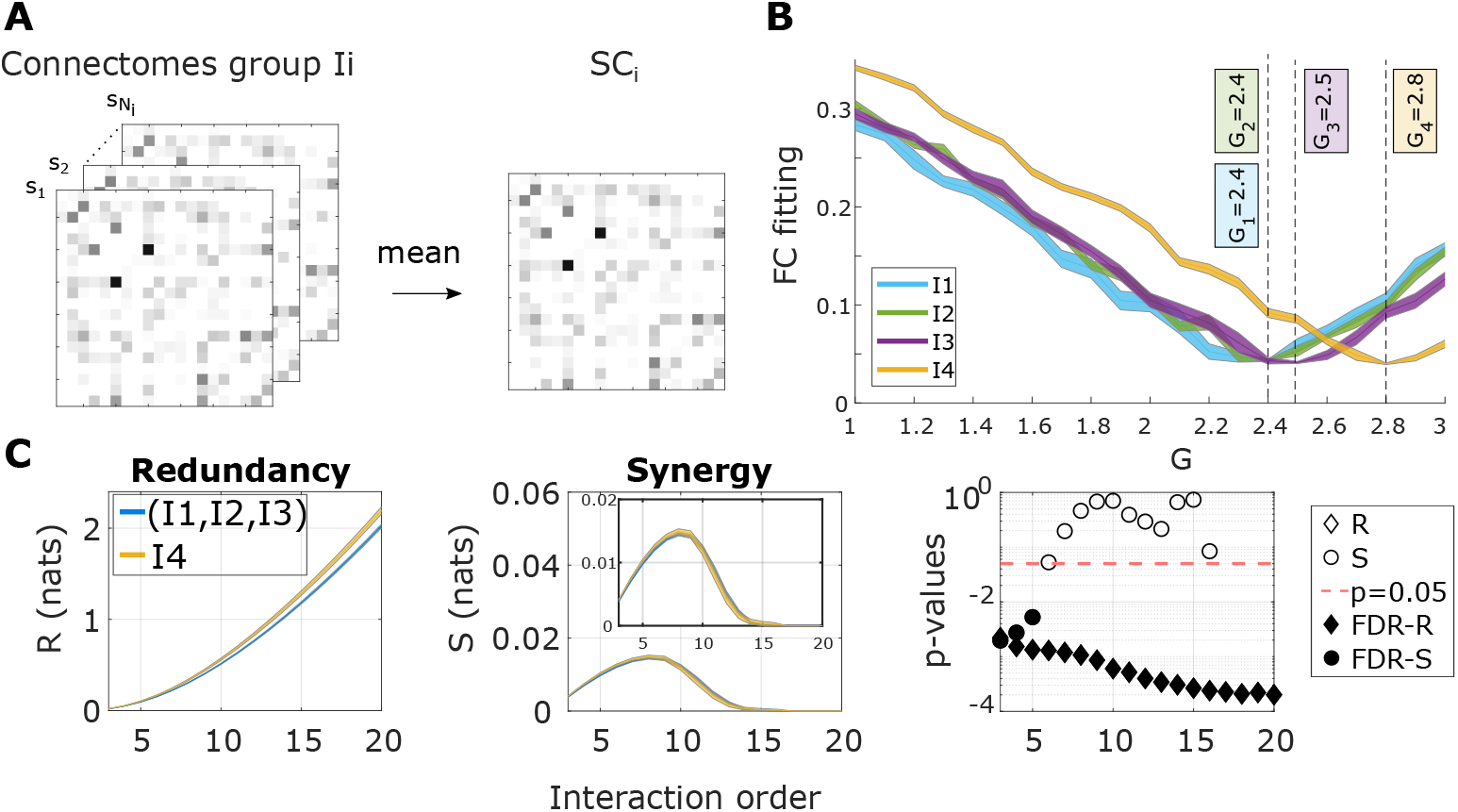
The DMF model reproduces the age variations in the high-order interactions of redundancy and synergy. **A**: First, the within age-group average SC matrix was calculated. **B**: Next, the optimal *G* value for each group was obtained. **C**: We simulated brain activity within each group using the average SC and the corresponding optimal *G_i_*, computed the O-information as a function of the interaction order, and separated sets of elements into the dominantly redundant (positive O-information values) and synergistic (negative O-information values). Here, the total redundancy (R) and synergy (S) was obtained as the average O-information over the redundant and synergistic sets, respectively. The p-values of the Wilcoxon test are also depicted as a function of the interaction order, after comparing the values in I4 versus the ones obtained from the combination of I1,I2, and I3. When the value of redundancy (or synergy) survived the false discovery rate correction, the diamonds (or circles) were filled.

We computed the O-information for all the subsets of brain regions of size 3 ≥ *n* ≥ 20, where *n* represents the interaction order. For each order, n-plets with positive and negative values of the O-information — called for simplicity ‘redundancy’ and ‘synergy,’ respectively — were calculated. Wilcoxon tests were performed to compare the average values of redundancy and synergy in the older participants (I4) with the values obtained from the groups of younger participants following previous work [24]. This is illustrated in Fig. 2C. The DMF model reproduced the age differences in redundancy and synergy reported in [24]. Moreover, the differences in redundancy between I4 and the rest of groups (I1,I2,I3) are statistically significant at all orders after the multiple comparison correction of the false discovery rate. Interestingly, although the DMF model was fitted only using pairwise FC values, the simulated dynamics captures similar profiles of high-order statistics than our fMRI data.

### A connectome-based model of brain ageing

Motivated by the fact that the DMF model (connected with the average SC within each age group) reproduced the high-order interaction patterns of redundancy and synergy very precisely, we asked whether varying the SC of the young population was sufficient to reproduce the high-order functional aspects in the older participants. We then studied the relationship between the weights of SC from the youngest group I1 and the corresponding weights from the oldest group I4 through a parabolic fitting (for further details see Methods). This second-order polynomial fitting revealed a non-linear reduction of the anatomical weights throughout the brain, in agreement with previous work [5,17,19] (Fig. 3A). Next, the fitted polynomial was used to simulate the effects of ageing in each of the young participants belonging to I1, thus resulting in an “aged” version of their connectomes (Fig. 3B). Each of these *synthetic* aged connectomes was used to run a set of simulations using the DMF with the optimal value G_4_. Finally, the high-order interactions of the simulated time-courses were calculated via separating the O-information into redundancy and synergy terms. Our results, illustrated in Fig. 3C, showed that the synthetically simulated aged participants also reproduced the functional changes observed empirically, exhibiting significant (FDR-corrected) increased redundancy at all orders.

**Fig 3.**
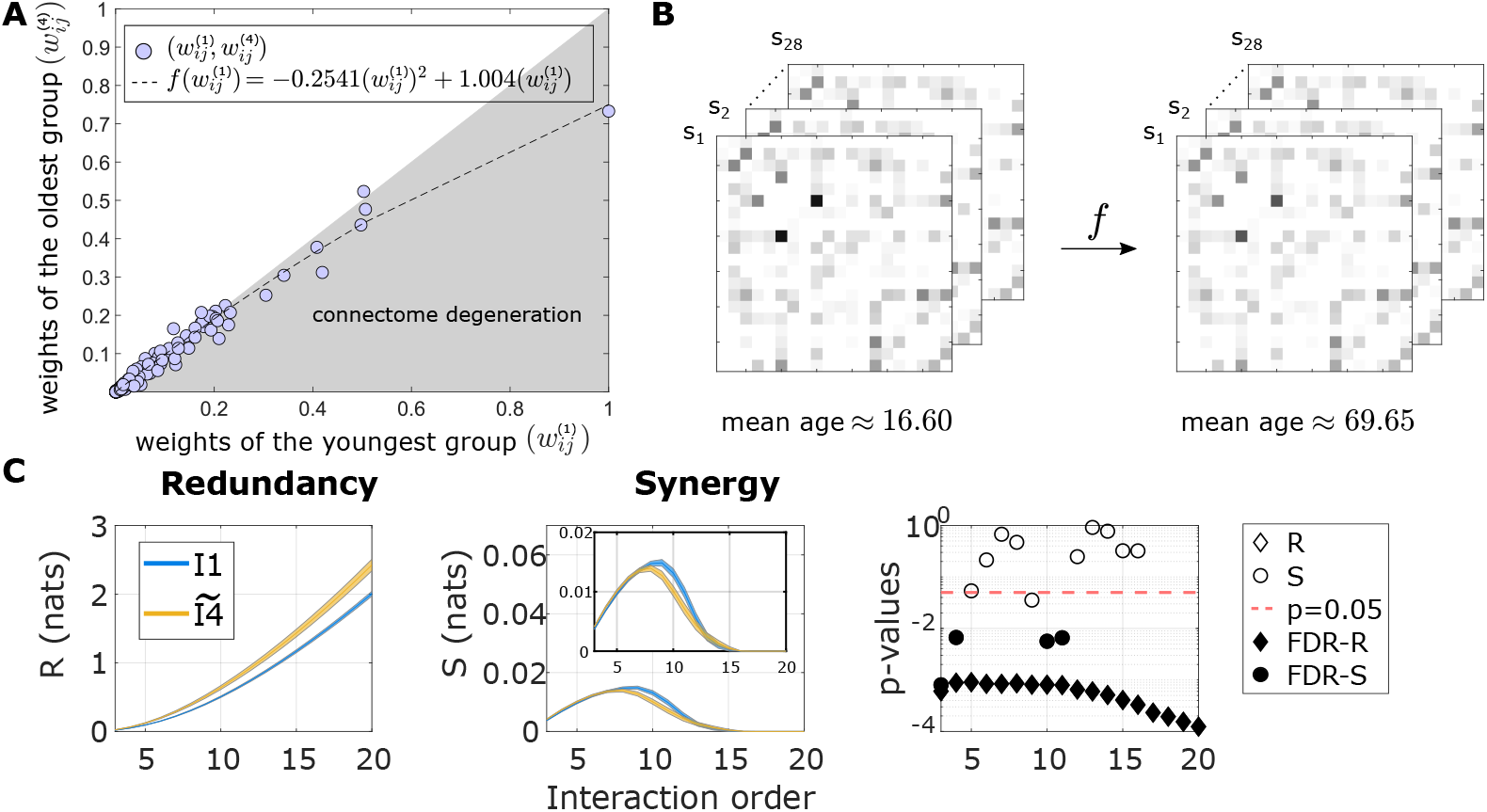
Connectome-based ageing model: **A:** A polynomial fit of second degree was used to link the weights of the average connectome within I1, denoted by 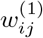, and the corresponding ones in I4 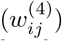. **B:** (left) Twenty-eight empirical young connectomes are transformed by the second-order polynomial fit, obtaining the synthetic aged connectomes (right). **C:** We simulated the DMF model of the aged connectomes and the optimal value *G*_4_. The O-information was assessed and separated into the total redundancy (left) and synergy (center). The third panel at the right corresponds to the p-values of the Wilcoxon test after comparing the redundancy and synergy of the synthetic I4 group with the ones in I1. When the value of redundancy or synergy survived multiple-comparison correction, both the diamonds and circles were filled.

To ensure that the good performance was not the result of simply taking *G*_4_ in the DMF modelling of the youngest group, the same analysis was repeated using a linear (instead of quadratic) model of connectome degreneration. The linear ageing model did not reproduce the observed redundancy variations between age groups, confirming that the non-linear heterogeneity in the age-related connectome degeneration is crucial to explain the observed changes in the high-order functional statistics. In the following section we dive deeper into the topological structure of these anatomical changes.

### Connectome degeneration heterogeneity revealed two differentiated communities of age-related links

We have shown that our connectome degeneration model based on a second-degree polynomial, reproduced the structure of high-order synergetic and redundant interactions of the oldest group. Interestingly, the non-linearity implies that not all the connectome links age in the same way. To further investigate this issue, we assessed across all participants in our cohort (N=161) the association between SC weights and age, by calculating the Pearson correlation *r* between age and each link of the SC matrix. This is illustrated in Fig. 4A.

**Fig 4.**
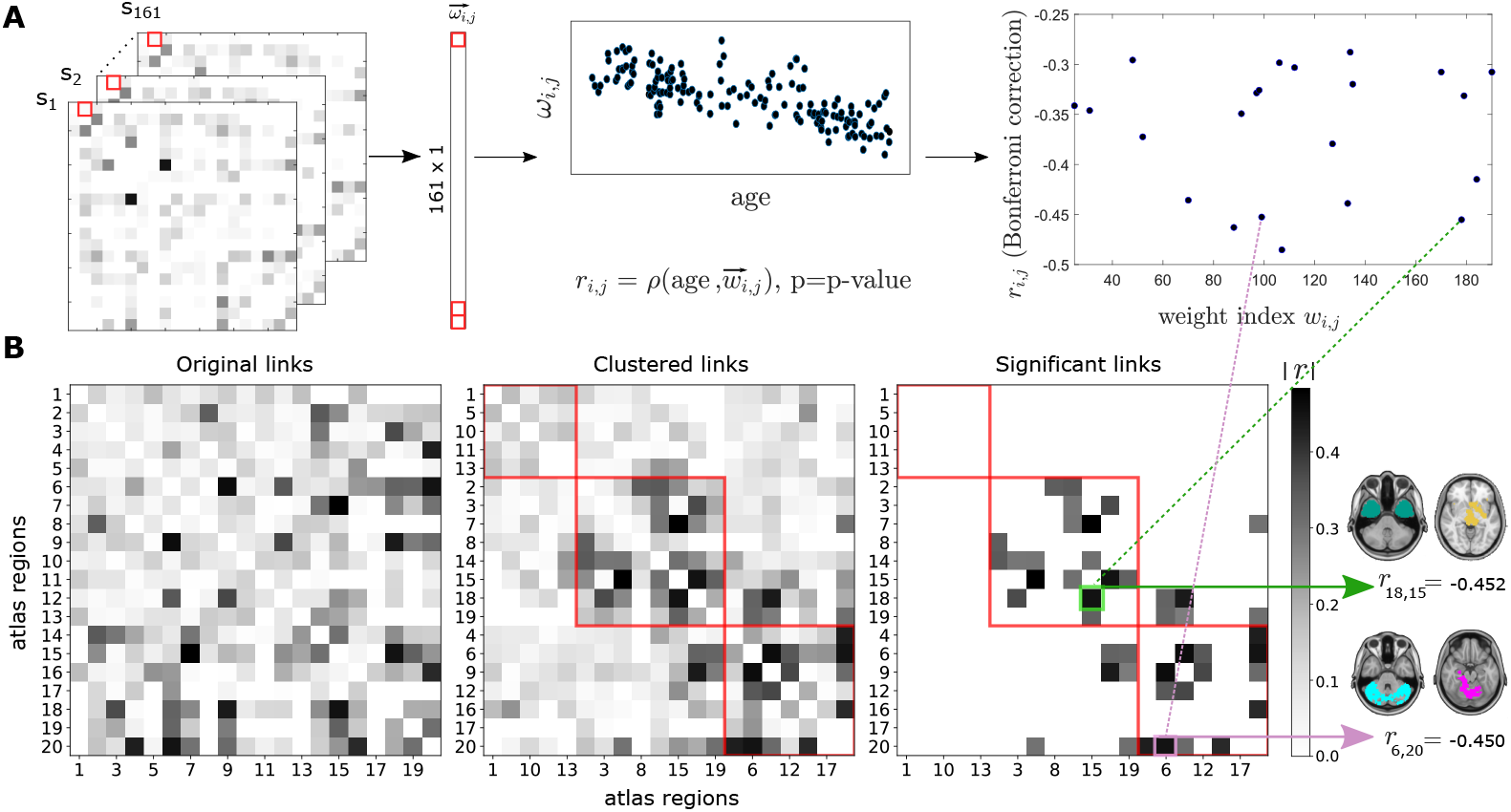
Heterogeneity of the connectome degeneration: **A:** The Pearson correlation *r* between age and individual weights *w_ij_* of the SC matrix was calculated across all the different participants (N=161). A correction for multiple comparisons was also applied. The final number of weights which survived to multiple comparisons is represented in the right panel, with values ranging from −0.25 to −0.50. **B:** We built a new connectivity matrix using as links the values of r obtained for each weight (left panel). After applying the Louvain Method of community detection to this matrix, three major communities were found (center), but only two communities had significant values of links (right). As an illustration, we showed one arbitrary link within each of these communities (colored in green and pink).

By using the correlation values as links of a new matrix (Fig. 4B), we applied the Louvain community detection method and obtained three distinct communities that aged differently. Moreover, after correcting for multiple comparisons two communities contained significant values. All the survived correlations had *r* values ranging from −0.25 to −0.5, thus showing a reduction in SC when age progressed.

Of note, the second community was dominated by interactions involving the brain atlas regions 15 and 18, showing the highest values of node-strength of the matrix after multiple comparison correction. These regions encompass several subcortical structures, such as the striatum, thalamus, brain stem, amygdala and the hippocampus (for a complete description of all regions in the atlas, see references [51,52]). In contrast, the third community showed that regions 6,9 and 20 had higher values of strength, indicating a major contribution of these three regions. Importantly, a common structure present in these three regions is the cerebellum. Therefore, the two communities exhibited age-induced reduction in within-community correlations, but in one, the degeneration was centered around the connectivity of the striatum and hippocampus, and in the other around that of the cerebellum.

## Discussion

In this article we used a combination of functional and diffusion MRI data together with DMF whole-brain modelling to investigate the mechanisms underlying age-variations in the structure of high-order functional interactions. The DMF model successfully reproduced the increased redundancy-dominated interdependencies of BOLD activity across brain areas in the older participants, and across all interaction orders, in full agreement with recent observations Ref. [24]. Furthermore, we provided evidence that these high-order functional changes are driven by localised non-lineal processes of neurodegeneration in the connectome. Leveraging this finding, we proposed a non-linear connectome-based degeneration model of ageing, which can be applied to young connectomes to simulate age-induced changes in functional brain patterns.

Whole-brain models of neuronal activity have significantly increased our understanding of how functional brain states emerge from their underlying structural substrate, and have provided new mechanistic insights into how brain function is affected when other factors are altered such as neuromodulation [28, 53, 54], connectome disruption [38, 55], or external stimuli [56, 57]. Adding to these findings, the present results provide a causal link between a localised connectome-based degeneration model of aging and age-variations of high-order functional interdependencies. These results establish a first step towards explaining how the reconfiguration of brain activity along the lifespan intertwines with changes in the underlying neuroanatomy.

Our results revealed two communities with differentiated age-induced deteriorated connectivity, one focused on the striatum and hippocampus, and the other on the cerebellum. In relation to striatal connectivity, previous studies showed that the fronto-striato-thalamic circuit was the most dominant for age prediction in healthy participants [58]. Moreover, age-related deterioration of striatal connectivity has also been associated with reduced performance in rest [26] and action selection tasks [59], inhibitory control [60], and executive function [61]. In relation to hippocampus, a gold-standard structure affecting memory-impaired degenerative diseases, is also affected in normal aging [62], with implications in spatial and episodic memories processing [63]. In relation to cerebellar connectivity, both sensorimotor and cognitive task performance in the older population has been shown to be associated with cerebellar engagement with the default mode network and striatal pathways [64]. The connectivity between cerebellum and striatum was also shown to be affected by age and exhibited relations with motor and cognitive performance [65]. Therefore, our results provide further support for the important behavioral implications that age-disconnection has on these circuits.

### Limitations and future work

This work makes use of a brain parcellation of only 20 regions from the brain hierarchical atlas (BHA) [51]. It has to be noted that, compared to other parcellations that focus into the cerebral cortex, the BHA encompasses the whole brain including brainstem, cerebellum, thalamus, striatum, amygdala, hippocampus, and cerebral cortex. While this parcellation was shown to maximize the cross modularity index between the functional and structural data, future work may also consider other brain parcellations to elucidate the robustness of our results by studying if age related changes in SC can also explain the differences high-order functional interactions in whole brain models. Analogously, some variations in the MRI preprocessing pipeline could also affects our results [66], as previous works have shown that affect pairwise FC studies [67].

Our analyses assessing high-order functional interactions are based on some specific metrics such as mean values of O-information at each interaction order. Future studies may consider different algebraic or topological properties of the full O-information hypergraph [68–73], which may provide complementary insights. It is also worth noting that the reported values of the O-information are not indicative of ‘pure’ synergy or redundancy, but correspond to the balance between them. The O-information was chosen because it is a convenient measure to assess high-order effects up to relatively high orders. However, the O-information is a whole-minus-sum type of measure, and hence its analysis does not fully discriminate e.g. net increases in redundancy from decreases in synergy. Future studies could perform more detailed analyses by employing partial information decomposition (PID) measures [74–81].

### Final remarks

In summary, our results extend previous findings on high-order interdependencies of the ageing brain using a novel framework that incorporated whole brain modelling and connectome datasets along the lifespan. Whole-brain models have enhanced our understanding of the brain across different conditions, and therefore provide a highly promising avenue of research in the field of ageing neuroscience. In this context, our work constitutes the first step towards mechanistic explanations on how functional high-order interdependencies in the human brain are affected by the age-connnectome degeneration. Future work should validate our modelling approach in the presence of other forms of brain degeneration, such as the interplay between aging and pathologies like Alzheimer or Parkinson diseases.

## Materials and methods

### Participants

A cohort of *N* = 161 healthy volunteers with an age ranging from 10 to 80 years (mean age 44.35 years, SD 22.14 years) were recruited in the vicinity of Leuven and Hasselt (Belgium) from the general population by advertisements on websites, announcements at meetings and provision of flyers at visits of organizations, and public gatherings (PI: Stephan Swinnen). Informed consent was obtained before participation in the study, according to the local ethical committee for biomedical research and the Declaration of Helsinki. None of the participants had a history of ophthalmological, neurological, psychiatric, or cardio-vascular diseases potentially influencing imaging measures. The participants were divided into four distinct age groups: I1 consists of *N*_1_ = 28 participants with ages ranging from 10–20 years, I2 of *N*_2_ = 46 from 20–40 years, I3 of *N*_3_ = 29 from 40–60 years and I4 of *N*_4_ = 58 from 60-80 years.

### Image acquisition and preprocessing

Image acquisition was performed on a MRI Siemens 3T MAGNETOM Trio MRI scanner with a 12-channel matrix head coil. The anatomical images were acquired with a 3D magnetization prepared rapid acquisition gradient echo (MPRAGE) and the following parameters: repetition time (TR) = 2,300 ms,echo time (TE) = 2.98 ms, voxel size = 1 × 1 × 1.1 mm^3^, slice thickness =1.1 mm, field of view (FOV) = 256 × 240 mm^2^, 160 contiguous sagittal slices covering the entire brain and brainstem. The anatomical images were then used for preprocessing of the functional data, here acquired with a gradient echo-planar imaging sequence over a 10 min session using the following parameters: 200 whole-brain volumes with TR/TE = 3000/30 ms, flip angle = 90, inter-slice gap = 0.28 mm, voxel size =2.5×3×2.5 mm^3^, 80×80 matrix, slice thickness = 2.8 mm, 50 oblique axial slices, interleaved in descending order. Functional imaging preprocessing was performed following a similar procedure to that in Ref. [25]. The preprocessing pipeline included slice-time correction, head motion artifacts removal, intensity normalization, regressing out of the average cerebrospinal fluid and average white matter signal, bandpass filtering between 0.01 and 0.08 Hz, spatial normalization to a template of voxel size of 3 × 3 × 3 mm^3^, spatial smoothing, and scrubbing. This resulted in a total of 2514 time series of fMRI BOLD signal for each participant, corresponding to the functional partition used in Ref. [51]. Moreover, because for the calculation of high-order interactions at order *n* we have to deal with *n*–plets of region combinations (for details see the following subsections), we reduced complexity grouping the original 2514 regions into 20 final brain atlas regions, simply by averaging the time series of all regions within a given atlas region. For this stage, we made use of the Brain Hierarchical Atlas [51], that has been previously used [82–85]. The partition of 20 regions is the one that maximized the cross-modularity, a metric that accounts for the triple optimization of the functional modularity, the structural modularity and the similarity between structural and functional regions (for details see Ref. [51]). To obtain the structural connectivity matrices, we acquired diffusion weighted single shot spin-echo echo-planar imaging (DTI SE-EPI) images with the following parameters: TR = 8,000 ms, TE = 91 ms, voxel size = 2.2 × 2.2 × 2.2 mm^3^, slice thickness = 2.2 mm, FOV = 212 × 212 mm^2^, for each image, 60 contiguous sagittal slices were acquired covering the entire brain and brainstem. A total number of 64 volumes were acquired corresponding to different gradient directions with b=1000 s/mm^2^. One extra 3D diffusion image was acquired for b = 0 s/mm^2^, needed for the diffusion imaging preprocessing. Although full details are given in Ref. [58], the pipeline consisted in eddy current correction, motion correction, tensor estimation per voxel, fiber assignment, and functional partition projection to the individual diffusion space. This resulted in SC matrices of dimension 2514 × 2514, one per participant, and each matrix entry corresponding with the number of white matter streamlines connecting that given region pair. Finally, we reduced complexity of these matrices by grouping the 2514 × 2514 matrix into 20 ×20 using the BHA, and averaging the BOLD signals of all regions within a given atlas region.

### Whole-brain dynamic mean field model

To simulate neuronal activity of each region, we used Dynamic Mean Field Modelling (DMF) [28,31]. Each brain region is modelled by interacting neural inhibitory (I) and excitatory (E) populations. DMF assumes that the the inhibitory currents *I*^(I)^ are mediated by GABA-A receptors, and the excitatory ones *I*^(E)^ by NMDA receptors. The connectivity between two different nodes *n* and *p* is given by the *C_np_*. In this work, we took *C_np_* equal to the element of the structural matrix *SC_np_*. Summarizing, neuronal activity followed:

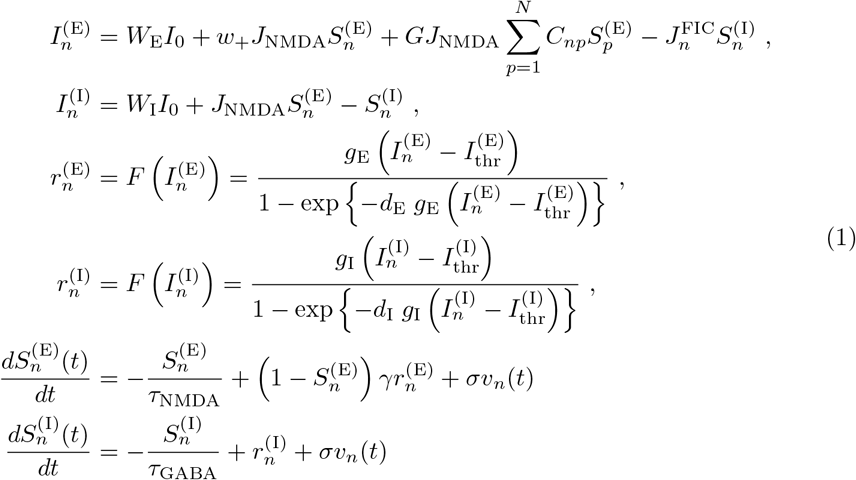

where the synaptic gating variable of excitatory pools is denoted by 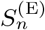 and the one for inhibitory populations as 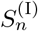. The excitatory and inhibitory firing rates are denoted by 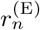 and 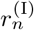 respectively. The feedback inhibitory control weight, 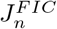 was adjusted for each node *n* in a way such that the firing rate of the excitatory pools 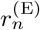 remains fixed at about 3 Hz, using the linear fitting strategy proposed by Herzog and colleagues [38]. The precise values of the parameters used here are given in Table 1.

**Table 1.** Dynamic Mean Field (DMF) model parameters

| Symbol | Parameter name | Value |
| --- | --- | --- |
| $I_0$ | External current | 0.382 nA |
| $W_E$ | Excitatory scaling factor for $I_0$ | 1 |
| $W_I$ | Inhibitory scaling factor for $I_0$ | 0.7 |
| $w_+$ | Local excitatory recurrence | 1.4 |
| $J_{\text{NMDA}}$ | Excitatory synaptic coupling | 0.15 nA |
| $I_{\text{thr}}^{(E)}$ | Threshold for $F(I_n^{(E)})$ | 0.403 nA |
| $I_{\text{thr}}^{(I)}$ | Threshold for $F(I_n^{(I)})$ | 0.288 nA |
| $g_E$ | Gain factor of $F(I_n^{(E)})$ | 310 nC<sup>-1</sup> |
| $g_I$ | Gain factor of $F(I_n^{(I)})$ | 615 nC<sup>-1</sup> |
| $d_E$ | Shape of $F(I_n^{(E)})$ around $I_{\text{thr}}^{(E)}$ | 0.16 s |
| $d_I$ | Shape of $F(I_n^{(I)})$ around $I_{\text{thr}}^{(I)}$ | 0.087 s |
| $\gamma$ | Excitatory kinetic parameter | 0.641 |
| $\sigma$ | Amplitude of uncorrelated Gaussian noise $v_n$ | 0.01 nA |
| $\tau_{\text{NMDA}}$ | Time constant of NMDA | 100 ms |
| $\tau_{\text{GABA}}$ | Time constant of GABA | 10 ms |

The complete DMF implementation was performed in Matlab, and is freely available at https://gitlab.com/concog/fastdmf.

#### Haemodynamic model

The excitatory firing rates 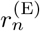 were transformed into BOLD-like signals, following a well-known hemodynamic model [86]. It is assumed that an increment in the firing rate 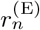 triggers a vasodilatory response *s_n_*, that produces a blood inflow *f_n_*, and changes the blood volume *v_n_* and the deoxyhemoglobin content *q_n_*. In particular, we modeled these interactions as:

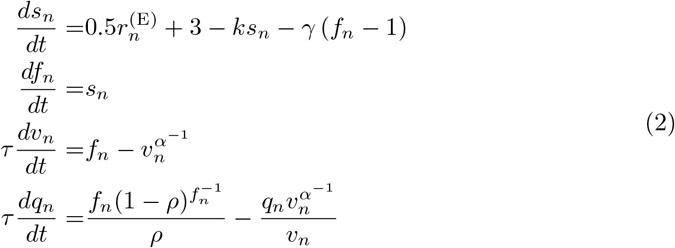

where *ρ* is the resting oxygen extraction fraction, *τ* is a time constant and *α* represents the resistance of the veins. The BOLD-like signal of node *n*, denoted *B_n_*(*t*), is a non-linear function of *q_n_*(*t*) (deoxyhemoglobin content) and *v_n_*(*t*) (blood volume), that can be written as:

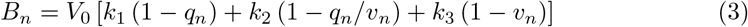

where *V*_0_ represent the fraction of venous blood (deoxygenated) in the resting-state, and *k*_1_ = 2.77, *k*_2_ = 0.2, *k*_3_ = 0.5 are kinetic constants, chosen from [86].

The numerical integration of the system in Eq. (2) was performed using Euler method, using an integration step of 1 ms. The signals were finally band-pass filtered between 0.01 and 0.1 Hz with a 3^rd^-order Bessel filter. To match the duration of the BOLD signals obtained from the participants of this study, we simulate 160 time-points of BOLD signals corresponding to 8 minutes for each brain region.These BOLD-like signals were the ones used for the calculation of the FC matrices and the O-information, the latter used for the calculation of high-order synergistic and redundant interactions.

### Model fitting across age groups

We ran the DMF model using the average connectome *SC_i_* per age group, and chose the corresponding global coupling parameter *G_i_* by minimizing the Kolmogorov-Smirnov distance between the two distributions of FC, one obtained from simulations and the other corresponding to the empirical FC (the one obtained from the real functional data). In both cases, we built FC matrices by calculating the mutual information between pairs of time series using Gaussian Copulas. For choosing the minimizing G, we varied it from 1 to 3 with steps of size 0.1. For each *G* value, we run 112 simulations using different random seeds. We obtained a convex curve where the x-axis represents the *G* values and the y-axis is the Kolmogorov–Smirnov distance. The value of *G* corresponding to the minimum Kolmogorov-Smirnov distance between the real and simulated data represents the optimal model.

### Connectome-based ageing model

We analysed the dependencies between the average SC in group I4 vs the group I1. The best fitting was achieved by a second-degree polynomial *f*, which was used as the connectome-based model for aging. For each participant of the youngest group, we modeled a synthetic *aged* version of his connectome. To obtain this synthetic connectome, we applied *f* for each element of the matrix SC of each participant within the group I1. Finally, we used these synthetic anatomical connectivity matrices and the parameter *G*_4_ = 2.8 as inputs for the DMF. For each synthetic SC, we run a number of four different simulations and obtained for each one a total number of 112 whole-brain simulations, corresponding to different initial conditions in the simulations.

### Communities of age-related links

We first assessed the association for each individual connection and age. To do so, we calculated the Pearson correlation between each entry of the SC matrix and age, using the different participants as observations (N=161). To find different communities of age-related brain links, we made use of the Louvain community detection algorithm available in the Brain Connectivity Toolbox [87]. The optimal network partition corresponds to a subdivision of non-overlapping node groups (or communities) that maximize the within-group links and minimize between-group links. The network nodes were the 20 brain regions of the Brain Hierarchical Atlas [51], and the links were the absolute value of the Pearson correlation coefficients. After applying the community detection algorithm, we pruned the non-significant links using a Bonferroni correction.

### High-order functional interactions

Following Rosas *et al*. [88], we summarize here the main information-theoretic measures employed in this article. To begin, we defined the *total correlation* [89] TC and the *dual total correlation* [90] DTC as:

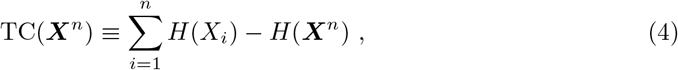

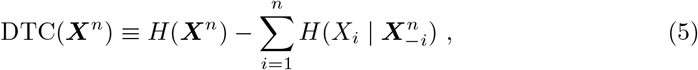

where *H*(·) represents the Shannon entropy, and 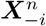 represents the vector of *n* − 1 variables composed by all vector components except *X_i_* i.e., (*X*_1_,…, *X*_*i*−1_, *X*_*i*+1_,…, *X_n_*). Both TC and DTC are non-negative generalizations of mutual information, meaning they are zero if and only if all variables *X*_1_,…, *X_n_* are statistically independent of one another.

For a set of *n* random variables ***X**^n^* = (*X*_1_,…, *X_n_*), the O-information [88] (denoted by Ω) was calculated as follows:

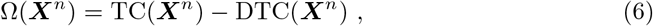

The O-information is a real-valued measure that captures the balance between redundancies and synergies in arbitrary sets of variables, thus extending the properties of the interaction information of three variables [91] to larger sets (see related discussion in Ref. [48]). In particular, the sign of the O-information serves to discriminate between redundant and synergistic groups of random variables: Ω > 0 corresponds to redundancy-dominated interdependencies, while Ω < 0 characterizes synergy-dominated variables.

From the time series of *n* different brain regions, we built vectors ***X**^n^* and computed all these quantities using Gaussian Copulas [92]. This approach exploits the fact that the Mutual Information does not depend on the marginal distributions, and therefore, the different quantities can be conveniently transformed to Gaussian random variables from which efficient parametric estimates of high-order interactions exist. All these quantities are always computed using natural logarithms.

### Interaction order, redundancy and synergy

We calculated the O-information for all the different *n*–plets of brain regions, with 3 ≥ n ≥ 20. Per participant, we computed the average O-information in which the region *m* participates when interacting with other *n* regions as

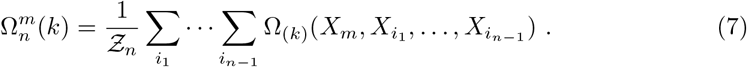

Above, *k* is the participant’s index, *m* the interacting region index, *n* the interaction order, and

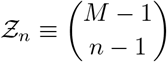

is the total number of subsets of size *n* – 1 in a brain partition of *M* regions (in this work we used *M* = 20). The summations in Eq. (7) included all the n-plets that included *X_m_*. Finally, the grand average O-information of order *n* is calculated averaging over all *M* regions.

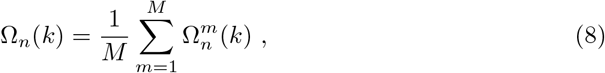

We then split the O-information on positive and negative values using

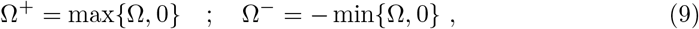

so that Ω = Ω^+^ – Ω^−^. Using these quantities, we calculated the following metrics for redundancy and synergy, for each subject *k*, interacting region *m*, and interaction order *n*:

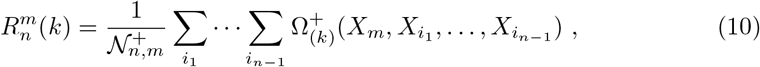

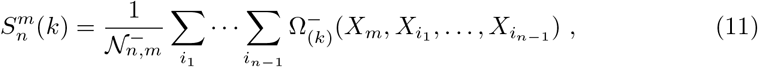

where 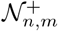 and 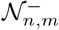 represent the number of n-plets with positive and negative O-information values, respectively.

Finally, the average of these quantities over all subjects and regions can be also calculated as

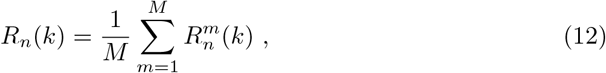

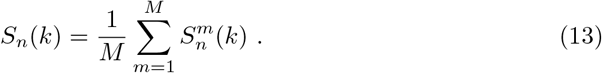

The code to compute all these metrics is available at https://github.com/brincolab/High-Order-interactions.

## Acknowledgments

M.G. was partially supported by CONICYT-PFCHA/ Doctorado Nacional/2019-21190577. R.C. acknowledges financial support from Fondecyt Iniciación 2018 Proyecto 11181072. P.M. was funded by the Wellcome Trust (grant no. 210920/Z/18/Z). F.R. acknowledges the support of the Ad Astra Chandaria Foundation. P.O. is funded by Fondecyt Regular grant 1211750 and ANID-Basal Project FB0008. J.M.C acknowledges financial support from Ikerbasque (The Basque Foundation for Science) and from the Ministerio Economia, Industria y Competitividad (Spain) and FEDER (grant DPI2016-79874-R), as well as from the Department of Economic Development and Infrastructures of the Basque Country (Elkartek Program, KK-2018/00032, KK-2018/00090, and KK-2021/00009). The Centro Interdisciplinario de Neurociencia de Valparaíso (CINV) is a Millennium Institute supported by Grant ICN09-022 (ICM-ANID).

